# Asymmetry of an intracellular scaffold at vertebrate electrical synapses

**DOI:** 10.1101/173955

**Authors:** Audrey J Marsh, Jennifer Carlisle Michel, Anisha P Adke, Emily L Heckman, Adam C Miller

## Abstract

Neuronal synaptic connections are electrical or chemical and together are essential to dynamically defining neural circuit function. While chemical synapses are well known for their biochemical complexity, electrical synapses are often viewed as comprised solely of neuronal gap junction channels that allow direct ionic and metabolic communication. However, associated with the gap junction channels are structures observed by electron microscopy called the Electrical Synapse Density (ESD). The ESD has been suggested to be critical for the formation and function of the electrical synapse, yet the biochemical makeup of these structures is poorly understood. Here we find that electrical synapse formation *in vivo* requires an intracellular scaffold called Tight Junction Protein 1b (Tjp1b). Tjp1b is localized to electrical synapses where it is required for the stabilization of the gap junction channels and for electrical synapse function. Strikingly, we find that Tjp1b protein localizes and functions asymmetrically, exclusively on the postsynaptic side of the synapse. Our findings support a novel model in which there is molecular asymmetry at the level of the intracellular scaffold that is required for building the electrical synapse. ESD molecular asymmetries may be a fundamental motif of all nervous systems and could support functional asymmetry at the electrical synapse.

## Paper

Fast communication between neurons in the brain is achieved by chemical and electrical synapses. Both synapse types are specialized neuronal adhesions allowing for communication, with each comprised of unique biochemical complexes. At chemical synapses, presynaptic active zones are specialized for neurotransmitter release and appose the postsynaptic density (PSD), which contains a set of intracellular scaffolds required for the trafficking, stability, and function of neurotransmitter receptors^1,2^. By contrast, electrical synapses are composed of neuronal gap junction (GJ) channels that allow direct cytoplasmic continuity between neurons. The GJs are composed of two hemichannels contributed by pre- and postsynaptic neurons, with each hemichannel comprising a hexamer of Connexin (Cx) proteins in vertebrates^3,4^. While many electrical synapses allow ionic flow equally in either direction^3,4^, others are rectified with flow being biased in one direction over the other^4–6^. Such functional asymmetries can occur in association with molecular asymmetries found at the level of the channels, with hemichannels created from different GJ-forming proteins contributed from each side of the synapse^6–9^. While much attention has been paid to neuronal GJ channel composition, little is known about scaffolding proteins localized to these connections. Electron microscopy of electrical synapses characteristically exhibits electron dense structures adjacent to the GJs on both the pre- and postsynaptic side of the synapse^10,11^. These electron dense structures, that we have termed the Electrical Synapse Density (ESD)^12^, likely represent macromolecular complexes associated with the GJ channels, and support their structure and function. However, the ESD proteins, their distributions, and functions are not well understood.

We use the electrical synapses of the Mauthner neural circuit in larval zebrafish, *Danio rerio* to investigate the molecular makeup of the ESD *in vivo* (Fig. 1A). The circuit drives a fast escape response to threatening stimuli^13^, with the two Mauthner neurons receiving input from sensory afferents and sending presynaptic axons down the length of the spinal cord where they activate a distributed postsynaptic network, including electrically activating the segmentally-repeated Commissural Local (CoLo) interneurons^9,14,15^. In a CRISPR-based reverse genetic screen we identified Tight Junction Protein 1b (Tjp1b) as being required for electrical synapse formation (Fig. 1B-E)^16^. Tjp1b is a highly-conserved homologue of mammalian Tjp1, also known as ZO-1, a multi-domain cytoplasmic scaffolding protein of the membrane-associated guanylate kinase (MAGUK) family (Fig. S1A). Tjp1/ZO-1 is best known for its role at epithelial tight/adherens junctions where it couples intercellular adhesion to intracellular actin cytoskeleton^17,18^. Within the brain it is localized at electrical synapses^19–21^, yet its function and fine-scale distribution remains unknown. In mammals, there are three related Tjp-encoding family-member genes, *tjp1, tjp2*, and *tjp3* – in zebrafish this family has expanded to *tjp1a, tjp1b, tjp2a, tjp2b*, and *tjp3* (Fig. S1A)^22^. We screened CRISPR-injected mosaic animals and found that of these five genes only *tjp1b* causes a defect in electrical synapse formation (Fig. S1B,C). To test whether the loss of *tjp1b* caused functional defects at electrical synapses we examined the effects on passage of the GJ-permeable dye neurobiotin (Nb) through M/CoLo synapses. We retrogradely labeled Mauthner axons with Nb from caudal spinal cord transections and detected Nb within the CoLo cell bodies (Fig. 1F-H)^9,15^. In mutants we found that Nb failed to transfer from Mauthner to CoLo (Fig. 1H-I). Taken together we conclude that the MAGUK scaffold Tjp1b is required for M/CoLo electrical synapse formation and function.

**Fig. 1.**
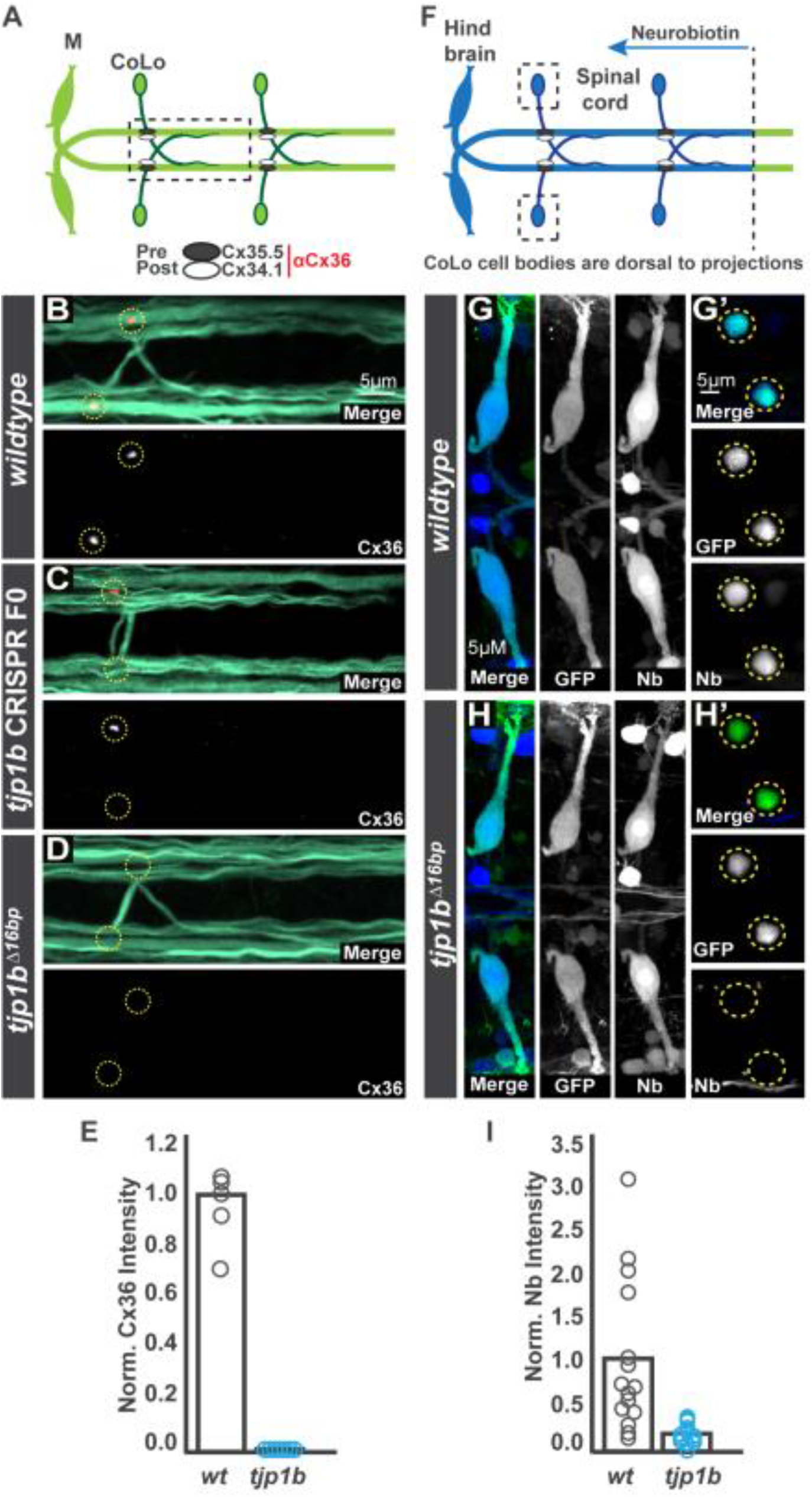
The MAGUK scaffold Tjp1b is required for electrical synapse formation and function. **A.** Schematic of the Mauthner circuit in dorsal view with anterior to the left (here and in all figures). Neurons and synapses of the hindbrain and two of thirty spinal segments are shown. Mauthner neurons send axons into the spinal cord making electrical synapses (black and white ovals) with CoLos. The M/CoLo synapses require two Cxs for formation, presynaptic Cx35.5 (black ovals) and postsynaptic Cx34.1 (white ovals). Boxed line denotes region of spinal cord shown in B-D. **B-D.** Dorsal views of a spinal cord segment from 5 days post fertilization zebrafish larvae. Images are maximum intensity projections of ~5μm (here and in all figures, unless otherwise noted) stained for neurofilaments (RMO44, green) and anti-human-Connexin36 (Cx36, red), which labels both Cx35.5 and Cx34.1^9^. Neighboring panels show individual Cx36 channels. **B.** *Wildtype* (*wt*). **C.** Mosaicly mutant CRISPR-injected F0 animals targeting the *tjp1b* locus. **D.** Homozygous *tjp1b* mutant. **E.** Quantitation of Cx36 fluorescent intensities in wildtype and *tjp1b* mutants. In this and all other graphs, each circle represents the measurement from an individual animal. **F.** Schematic of retrograde labeling of Mauthner axons with the gap junction permeable dye Neurobiotin (Nb) applied from a caudal transection (dotted line). Boxed region denotes the CoLo cell bodies shown in G’ and H’. **G,H.** Hindbrain (~15μm projection) and (**G’,H’**) spinal cord images at the level of the cell bodies stained for anti-GFP (green) and Nb (blue). **G,G’.** *wt M/CoLo:GFP* transgenic. **H,H’.** Homozygous *tjp1b* mutant in *M/CoLo:GFP* transgenic background. **I.** Quantitation of the ratio of Nb in CoLo to Mauthner cell bodies in wildtype and mutants.

We next examined whether Tjp1b localizes to Mauthner electrical synapses in the spinal cord. Five Cxs have been shown to create electrical synapses in the mammalian brain^4,12^. We previously found that Mauthner electrical synapses require two Cxs, Cx35.5 and Cx34.1, for their formation (Fig. 2A), and each Cx is required for the other to localize to the synapse^9^. Previous work has found that *tjp1b* mRNA is expressed throughout the zebrafish central nervous system^22^, including in the regions where Mauthner and CoLos reside. To examine protein localization we used an antibody against the human Tjp1 and found staining present broadly throughout the nervous system (Fig. 2A), but also specifically at Mauthner electrical synapses colocalized with Cx35.5 and Cx34.1 (Fig. 2A’). In *tjp1b* mutants much of the broad Tjp1 staining remains (Fig. 2B), however Tjp1 staining at the electrical synapse is lost along with Cx35.5 and Cx34.1 (Fig. 2B’,C). We expect that the Tjp1 antibody recognizes multiple Tjp proteins given the large *tjp-family* in zebrafish (Fig. S1A). Further supporting this idea we found that Tjp1 staining in *tjp1a* mutant animals results in a loss of neighboring, non-synaptic staining while synaptic staining remains (Fig. S2). We conclude that Tjp1b localizes to and is required for the stabilization of both Cx35.5 and Cx34.1 at electrical synapses.

**Fig. 2.**
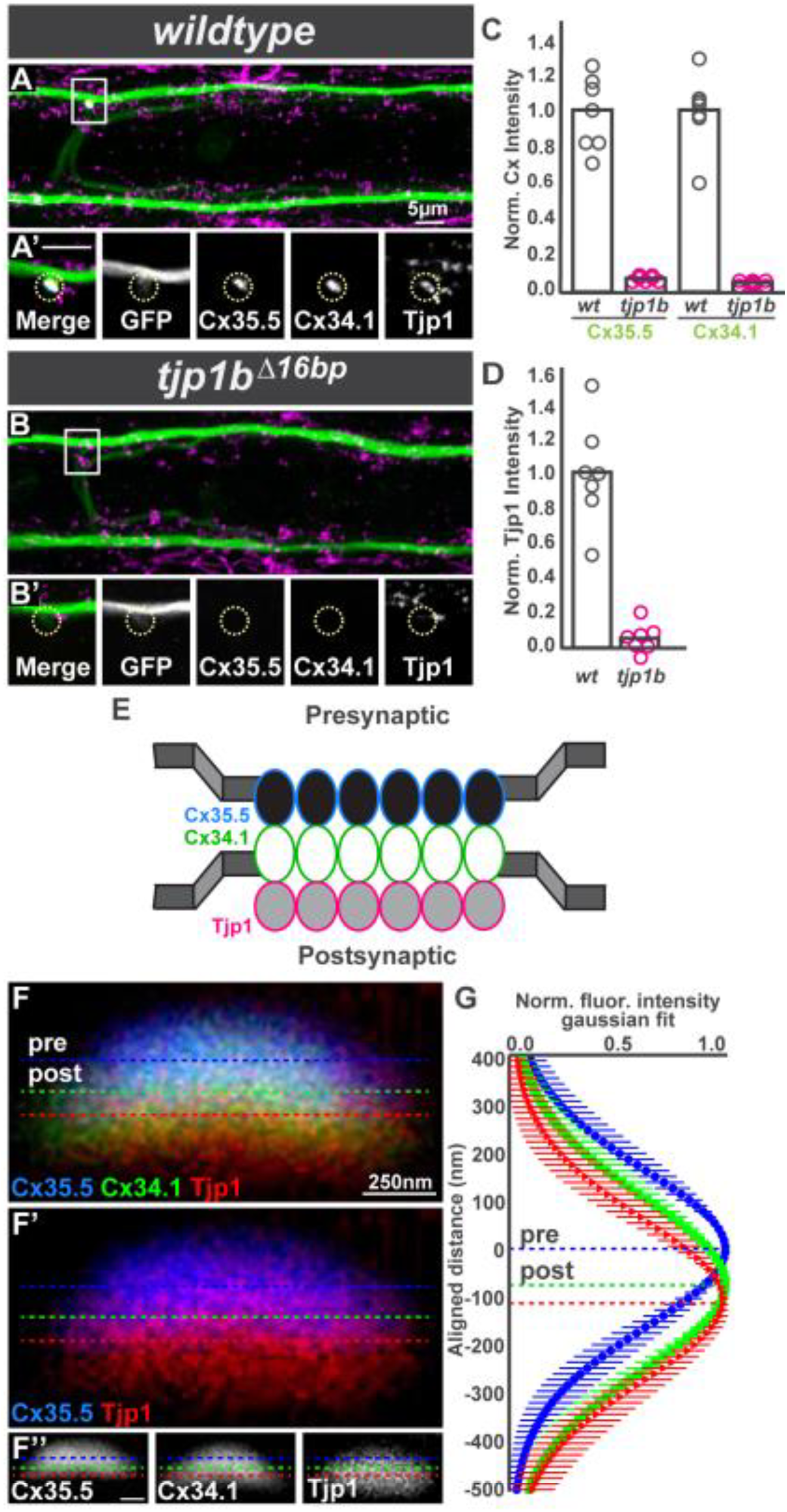
Tjp1b is asymmetrically localized at electrical synapses and required for pre- and postsynaptic Cx localization. **A,B.** Dorsal view projections from *M/CoLo:GFP* larvae stained for anti-GFP (green), anti-zebrafish-Cx35.5 (yellow), anti-zebrafish-Cx34.1 (cyan), and anti-human Tjp1/ZO1 (magenta). Boxes denote location of zooms. **A’,B’.** Individual Z-plane zooms. Yellow circle denotes location of M/CoLo site of contact. Individual channels are shown in neighboring panels. **A,A’.** *Wildtype* (*wt*) *M/CoLo:GFP* transgenic. **B,B’.** Homozygous *tjp1b* mutant in *M/CoLo:GFP* transgenic background. **C,D.** Quantitation of Cx35.5, Cx34.1 (**C**), and Tjp1 (**D**) fluorescent intensities in wildtype and *tjp1b* mutant animals. **E.** Model of protein localization observed at M/CoLo synapses. **F.** *wt* electrical synapse oriented with presynaptic Mauthner at the top and the postsynaptic CoLo at the bottom of the image (cells not shown). Larvae are stained for anti-zebrafish-Cx35.5 (blue), anti-zebrafish-Cx34.1 (green), and anti-human Tjp1 (red). Each dotted line represents the peak of fluorescence intensity of individual stainings for a line running vertically down the middle of the synapse. The Cx34.1 channel is removed in (**F’**) and individual channels are shown in (**F’’**). **G.** Fluorescent intensities from the Cx35.5, Cx34.1, and Tjp1 channels aligned to the center of the Cx35.5 peak fluorescence. Each point represents the average fluorescence intensity from 35 individual synapses in 8 animals at the specified distance away from the Cx35.5 peak fluorescence, with bars representing standard deviation. The peak-to-peak distance between the Cxs is ~81 nM, while Tjp1 staining is consistently ~33 nm further displaced away from Cx35.5.

We next investigated the precise localization, *i.e* pre- or postsynaptic, of the Cxs and Tjp1b at electrical synapses. Given that the mammalian Tjp1 binds to Cx36^19^ through highly-conserved domains present in the fish Tjp1b, Cx35.5, and Cx34.1 proteins (Fig. S1A)^9,22^, we hypothesized that Tjp1 staining would be associated with both the pre- and postsynaptic sides of the electrical synapse. Our previous work^9^ used genetic manipulations of Mauthner and CoLo neurons to show that the electrical synapses require Cxs asymmetrically – Cx35.5 is required presynaptically while Cx34.1 is required postsynaptically, creating a model molecularly asymmetric Cxs at the synapse (Fig. 2E). To examine whether we could observe the molecular asymmetry directly we used Cx35.5 and Cx34.1 specific antibodies^9^ and imaged M/CoLo synapses at high magnification. Synapses could be oriented towards the microscope lens *en face*, revealing a broad, curved surface (Fig. S3A), or could be observed on edge in cross-section (Fig. 2F, Fig S3B). We focused on the cross-section views as these gave the best orientation to view the pre- and postsynaptic localization of proteins. While both Cx35.5 and Cx34.1 are tightly localized to the synapse with overlapping tails of fluorescent distribution, the peak of staining for each is consistently offset from the other by ~81 nm (Fig. 2F,G; Fig. S3B-F). This supports our previous findings^9^ of molecular asymmetric Cxs. We next asked whether Tjp1 staining was symmetric or asymmetric and found the peak of Tjp1 staining was consistently offset by ~33 nm from the postsynaptic Cx34.1, and was ~118 nm away from the presynaptic Cx35.5 (Fig. 2F,G; Fig. S3B-F). These results support a model where Tjp1b is localized postsynaptically at these electrical synapses (Fig. 2E).

The apparent postsynaptic localization of Tjp1b suggests that it might function asymmetrically during electrical synapse formation. To test whether Tjp1b has a differential function in pre- and postsynaptic neurons, we created chimeric animals containing cells derived from two genetically distinct embryos by transplanting cells from GFP-expressing transgenic donors into non-transgenic hosts (Fig. 3A,B). In control chimeras with Mauthner or CoLo derived from *wildtype* transgenic donors we saw no effect on electrical synapse formation (Fig. 3C,D). By contrast, when only the postsynaptic neuron lacked *tjp1b* the Cxs failed to localize at electrical synapses (Fig. 3F,I). However, when the presynaptic Mauthner neuron lacked *tjp1b*, Cx localization was not affected (Fig. 3E,I). That is, Tjp1b function is required postsynaptically for both postsynaptic Cx34.1 and presynaptic Cx35.5 localization. Additionally, we note that Tjp1 staining is still present at the synapse when the presynaptic neuron is *tjp1b* mutant (Fig. 3E’), but not when the postsynaptic neurons are *tjp1b* mutant (Fig. 2F’), supporting the idea that the scaffold is localized and required postsynaptically. We conclude that Tjp1b is dispensable presynaptically, but localized and required postsynaptically for electrical synapse formation.

**Fig. 3.**
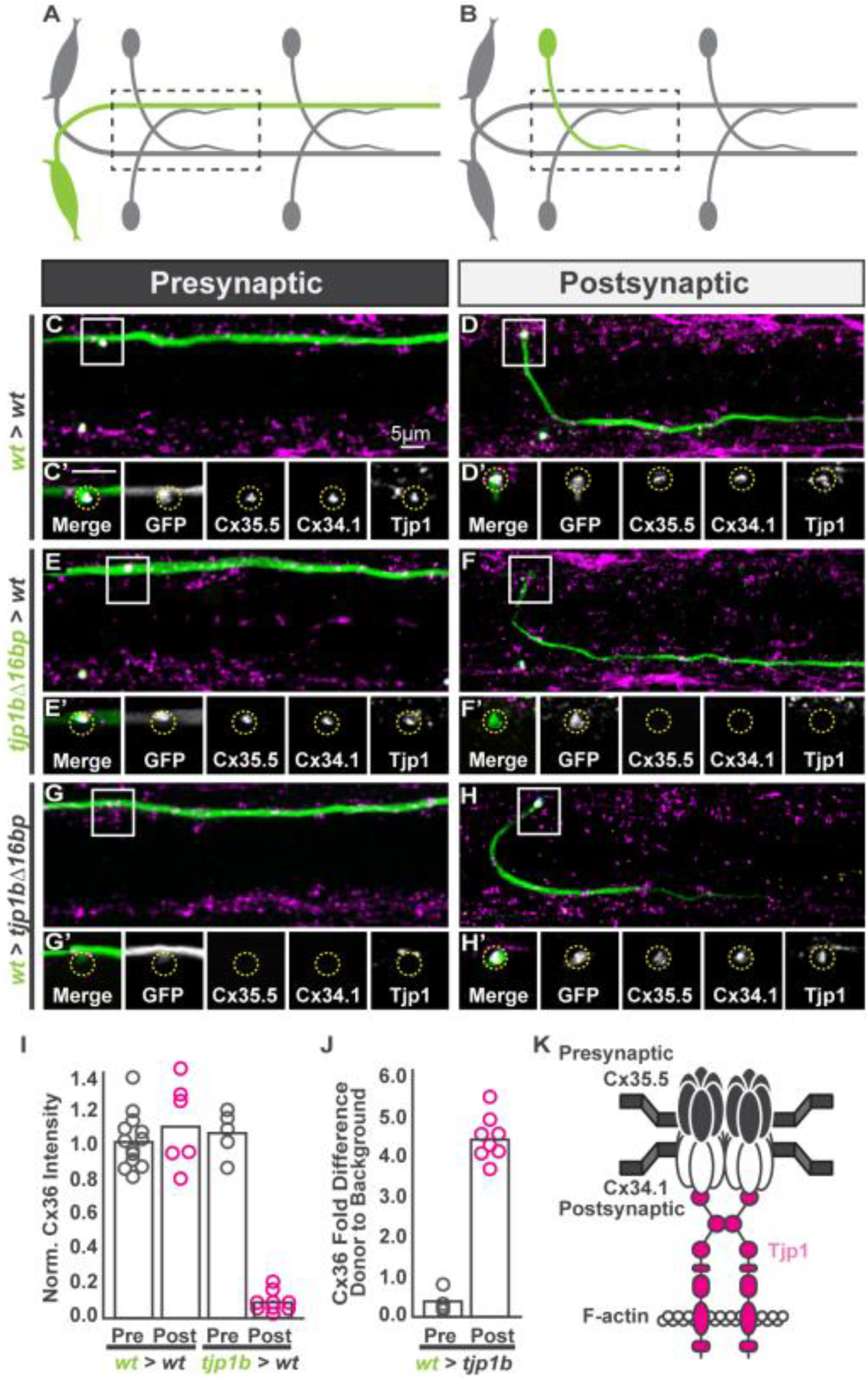
The exclusive asymmetric function of *tjp1b* postsynaptically is necessary and sufficient for electrical synapse formation. **A,B**. Schematics of chimeric larval circuits with the presynaptic Mauthner (**A**) or postsynaptic CoLo (**B**) derived from a transgenic donor. The genotype of the donor cell and host varies and is noted on the left of each set of images below. **C-H.** Dorsal views of chimeric larvae with GFP-marked cells (green) transplanted from a donor embryo into an unmarked host. Larvae are stained with anti-zebrafish-Cx35.5 (yellow), anti-zebrafish-Cx34.1 (cyan), and anti-human Tjp1/ZO1 (magenta). Boxes denotes location of single Z-plane zoom images in the ‘ panels. **C,D**. Control transplants from *wildtype* (*wt*) *M/CoLo:GFP* transgenic into a *wt* host. **E,F.** Transplants from *tjp1b* mutant *M/CoLo:GFP* transgenic into a *wt* host. **G,H.** Transplants from *wt M/CoLo:GFP* transgenic into a *tjp1b* mutant host. **I,J.** Quantitation of the ratio of Cx36 at donor-associated synapses to host synapses in chimeric embryos. **K.** Molecular model of postsynaptically localized Tjp1b required for asymmetric Cxs at the electrical synapse. Tjp1 may dimerize using its 2^nd^ PDZ domain^34^ and thereby cluster Cxs at the synapse. Molecular asymmetry may create functional asymmetry at these synapses.

Our results argue that Tjp1b is required for postsynaptic Cx34.1 hemichannel localization at synapses, which transsynaptically interacts with presynaptic Cx35.5 to stabilize the GJ channel. This is consistent with our previous findings that when Cx34.1 was removed postsynaptically, Cx35.5 could not stabilize presynaptically^9^. This would help explain why when we remove *tjp1b* postsynaptically we observe the loss of both postsynaptic Cx34.1 and presynaptic Cx35.5. This model predicts that upon *tjp1b* removal both Cx34.1 and Cx35.5 should still be expressed by their respective neurons, and synapse formation might be rescued if postsynaptic Tjp1b is resupplied. To test this we created chimeric animals as above except we transplanted from *wildtype M/CoLo:GFP* embryos into non-transgenic *tjp1b* mutant hosts. We found that a *wildtype* presynaptic neuron in a *tjp1b* mutant did not rescue the electrical synapse (Fig. 3G,J). By contrast, when only the postsynaptic neuron was wildtype (*i.e.* has Tjp1b) in an otherwise mutant animal the synapse was fully rescued (Fig. 3H,J). Thus postsynaptic Tjp1b is sufficient to rescue both postsynaptic Cx34.1 and presynaptic Cx35.5. We conclude that electrical synapse formation requires the exclusively postsynaptic function of Tjp1b, which supports the transsynaptic stability of the asymmetric neuronal GJ channels.

Our results strongly support the notion of molecular asymmetry at the level of the intracellular MAGUK Tjp1b at electrical synapses *in vivo* (Fig. 3K). This asymmetry of scaffold localization and function supports the molecularly asymmetric neuronal GJ channels found at these synapses^9^. This configuration of asymmetry at the membrane GJ hemichannels and the intracellular ESD is analogous to that found in their chemical synapse cousins. At chemical synapses, MAGUK scaffolds, such as PSD-95, are used extensively to regulate the trafficking, insertion, and modulation of neurotransmitter receptors to control synaptic function^2^. Intriguingly, *in vivo* electrical synapses can be rectifying, *i.e.* have biased flow through the GJ channel, with the proposed mechanism being constrained to asymmetry in the GJ forming proteins^7,6^. It is intriguing to speculate that the molecularly asymmetric MAGUK scaffold identified here may provide a platform to create functional asymmetry at electrical synapses, perhaps by allowing differential phosphorylation of Cx hemichannels^23^ or by affecting the local intracellular milieu in a manner that differentially gates the Cxs^24^. Future work will unravel the functional effects of molecularly asymmetric scaffolding at electrical synapses. What other asymmetries might be present at electrical synapses? In addition to Tjp1b, we previously identified Neurobeachin, an autism-associated protein, as being required postsynaptically for electrical synapse formation^15^. In contrast to what we find here for Tjp1b, Neurobeachin is localized to post-Golgi transport vesicles within the cytoplasm, and work in mouse and cell culture has shown that it is required for vesicular transport of proteins to the membrane^25,26^. One intriguing possibility is that Neurobeachin interacts directly with a Tjp1b/Cx34.1 complex being transported within the postsynaptic neuron to ensure its appropriate trafficking, localization, and stability at electrical synapses. We also expect a presynaptic scaffold to be present at these synapses given that electron microscopy of related synapses in goldfish shows electron dense structures both pre- and postsynaptically^10^; currently the molecular identity of such a scaffold is unknown. Further investigations will continue to extend the idea of molecular and functional asymmetry at electrical synapses and we expect that these structures will be critical to the development, function, and plasticity of neural circuits in all organisms.

## Methods

### Fish, lines, and maintenance

All fish were maintained in the University of Oregon’s fish facility under an Institutional Animal Care and Use Committee (IACUC) approved protocol #12498. Zebrafish, *Danio rerio*, were bred and maintained as previously described^27^. Animal care was provided by the University of Oregon fish facility staff and veterinary care was provided by Dr. Kathy Snell, DVM. All fish used on this project were generated in the AB x Tübingen hybrid background. With the exception of CRISPR injected and transplant hosts embryos, all fish had the enhancer trap transgene *M/CoLo:GFP (zf206Et)* in the background^14^. The *tjp1b^Δ16bp^* (*fh448*) and tjp1a^*Δ*2bp^ (*fh463*) lines were generated using the CRISPR Cas9 system^28^ by targeting the first common exon of its three mRNA isoforms. All experiments were performed at 5 days post fertilization (dpf). Newly generated mutant lines were Sanger sequenced to verify genomic changes.

### Immunohistochemistry

Anesthetized larvae were fixed for 3 hours in 2% trichloroacetic (TCA) acid or for 1 hour in 4% paraformaldehyde (PFA). Fixed tissue was then washed in PBS + 0.5% TritonX100, followed by standard blocking and antibody incubations. Tissue was cleared step-wise in a 25%, 50%, 75% glycerol series and was dissected and mounted in ProLong Gold antifade reagent (ThermoFisher, P36930). Primary antibodies used were: chicken anti-GFP (abcam, ab13970, 1:200), rabbit anti-human-Cx36 (Invitrogen, 36-4600, 1:200), mouse anti-RMO44 (Life Technologies, 13-0500, 1:100), mouse IgG1 anti-Tjp1 (ThermoFisher, 339100, 1:200), rabbit anti-Cx35.5 (Fred Hutch Antibody Technology Facility, clone 12H5, 1:200), and mouse IgG2A anti-Cx34.1 (Fred Hutch Antibody Technology Facility, clone 5C10A, 1:200). All secondary antibodies were raised in goat (Life Technologies, conjugated with Alexa-405, -488, -555, -594, or -633 fluorophores, 1:500). Neurobiotin was detected using fluorescently-tagged streptavidin (Life Technologies, conjugated with Alexa-633 fluorophores, 1:750).

### Confocal imaging and analysis

All images were acquired on a Leica SP8 Confocal using a 405-diode laser and a white light laser set to 491, 553, 598, and 639 nm depending on the dye. Each line’s data was collected sequentially using custom detection filters based on the dye. Most images were collected using a 40x, 1.2 numerical aperture (NA), water immersion lens. For each set of images the optical section thickness was calculated by the Leica software based on the pinhole, emission wavelengths, and NA of the lens. Images were processed and analyzed using FiJi^29^ software. Within each experiment all animals were stained together with the same antibody mix, processed at the same time, and all confocal settings (laser power, scan speed, gain, offset, objective, zoom) were identical. To quantify the presence or absence of staining at the synapse, a standard region of interest (ROI) surrounding each M/CoLo site of contact was drawn and the mean fluorescent intensity was measured. For neurobiotin backfills fluorescent intensity was measured using a standard ROI encompassing the entire Mauthner or CoLo cell body. For relative localization of Cx35.5, Cx34.1, and Tjp1 at synapses images were taken with a 63X, 1.4 NA, oil immersion lens at 5X digital magnification using a 1024^2^ pixel image. Each synapse was visually inspected for its orientation (see text and Fig. S3) and those with a view from the side were selected for protein localization analysis. The fluorescence intensity of each protein was analyzed at the middle-plane of the synapse, with a 1.2 μm ROI line drawn orthogonally from pre- to postsynaptic side of the synapse. For chromatic aberration controls, rabbit anti-Cx35.5 was stained with anti-rabbit-488, anti-rabbit-555, and anti-rabbit-633. For the localization of each protein, rabbit anti-Cx35.5, mouse IgG2A anti-Cx34.1, and mouse IgG1 anti-Tjp1 were stained with anti-rabbit-488, anti-mouse-IgG2A-555, and anti-mouse-IgG1-633. The fluorescence intensity of each individual channel was measured along the ROI and fit to a Gaussian curve. Each Gaussian curve was aligned based on the 488-channel and distances between the peaks of fluorescence were measured. Statistics were computed using Prism software (GraphPad). Figure images were created using FiJi, Photoshop (Adobe), and Illustrator (Adobe).

### CRISPR screening

Single guide RNAs (sgRNAs) were designed using the CRISPRscan database (http://www.crisprscan.org)^30^ to target a conserved exon of each individual *tjp* gene. sgRNAs were synthesized as previously described^28^ and were combined with Cas9 protein (IDT 1074181) at 37C for 5 minutes in the presence of 300mM KCl, and 4mM HEPES pH7.5 before injection^31^. All experiments used a final concentration of 1600 pg/nl of Cas9 protein; those targeting individual *tjp* genes used a final concentration of 200 pg/nl of sgRNA, while multiplexed *tjp* family injections included a total of 100 pg/nl of mixed sgRNAs. A total of 1 nl of Cas9/sgRNA complex was injected at the one cell stage and 5 dpf larvae were fixed in TCA and processed for immunohistochemistry. Synapse loss was assessed by counting the number of Cx36 punctae lost per total number of M/CoLo contacts counted per fish. The CRISPR mutagenesis rate was determined by fragment analysis using the CRISPR-STAT method^32^.

CRISPR targets, with the PAM site underlined:

*tjp1a:* GGGTTTGGAATCGCCATCTC<u>AGG</u>
*tjp1b:* GTGGGCTTGAGGCTCGCTGG<u>GGG</u>
*tjp2a:* GGCTTTGGCCTAGCGGTGTC<u>TGG</u>
*tjp2b:* GTTTGGATTGTCCCGGCCGC<u>CGG</u>
*tjp3:* GATTCCCACCGGTCCCGCCA<u>TGG</u>

### Neurobiotin retrograde labeling

Anesthetized 5 dpf embryos were mounted in 1.4% agar and a caudal transection through the dorsal half of the embryo was made with an insect pin at somite 20. A second insect pin loaded with 5% neurobiotin (Nb) solution was quickly applied to the incision. Anesthetized animals were unmounted from the agar and allowed to rest for 5 hours to allow neurobiotin to pass from Mauthner into CoLos, and were then fixed in PFA and processed for immunohistochemistry. CoLo axons project posteriorly for a maximum of two segments; therefore measurements of Nb in CoLo were analyzed at least three segments away from the lesion site.

### Blastula cell transplants

Cell transplantation was performed at the high stage approximately 3.3 hours into zebrafish development using standard techniques^33^. For “mutant into wildtype” experiments, animals heterozygous for the *tjp1b^Δ16bp^* mutation in the M/CoLo:GFP background were crossed and random progeny were used to transplant into non-transgenic *wildtype* hosts. Donor embryos were kept separate and genotyped at 1 dpf to assess the genotype of transplanted cells. For “wildtype into mutant” transplants transgenic *M/CoLo:GFP wildtype* animals were crossed to use as donors, and non-transgenic, heterozygous *tjp1b^Δ16bp^* animals were crossed for hosts and hosts were genotyped. Approximately 20 cells were deposited ~5-10 cell diameters away from the margin with a single embryo donating to 3-5 hosts. 5 dpf larvae were fixed in TCA and processed for immunohistochemistry.

## Author Contributions

All authors contributed to experiments and data analysis. A.C.M. wrote the manuscript. All authors edited the manuscript.

## Statement of Competing Interests

None.

## Materials and Correspondence

Adam Miller

## Acknowledgements

We thank the University of Oregon Fish Facility for superb animal care. We appreciate the discussions and advice from all lab members and members of the Institute of Neuroscience and the Institute of Molecular Biology at the University of Oregon. Funding was provided by the National Institute of Health Pathway to Independence Award R00NS085035 and the University of Oregon to A.C.M.

---

**Fig. S1.**
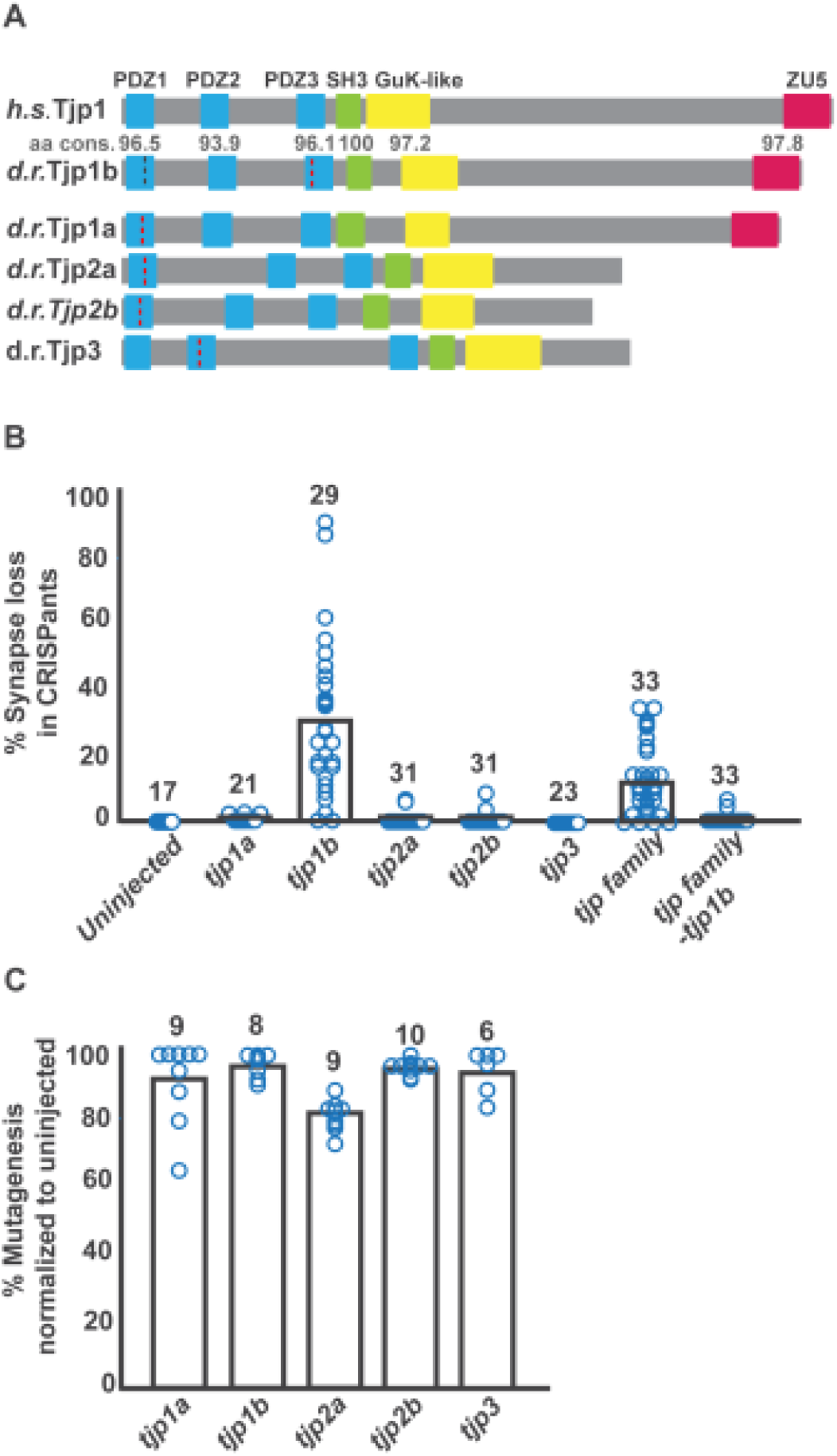
*tjp* gene family. **A.** Schematic of the primary structure of Tjp family proteins. Protein domains are depicted as colored boxes and the amino acid conservation (aa cons.) between zebrafish (*d.r.*) Tjp1b and human (*h.s.*) Tjp1 domains are noted. Black dotted line in Tjp1b represents location of CRISPR induced 16bp deletion (*fh448*) used throughout paper. Red dotted lines represent CRISPR target regions assayed in B and C. **B, C.** Quantitation of mosaic electrical synapse loss (**B**) and mutational efficiency (**C**) in CRISPR injected F0 embryos for each noted individual target or set of multiplexed targets. The “% Synapse loss” is the proportion of electrical synapses missing from at least 10, but on average 26, synapses sampled per animal. Mutagenesis was assessed by fragment analysis of PCR products at each targeted locus.

**Fig. S2.**
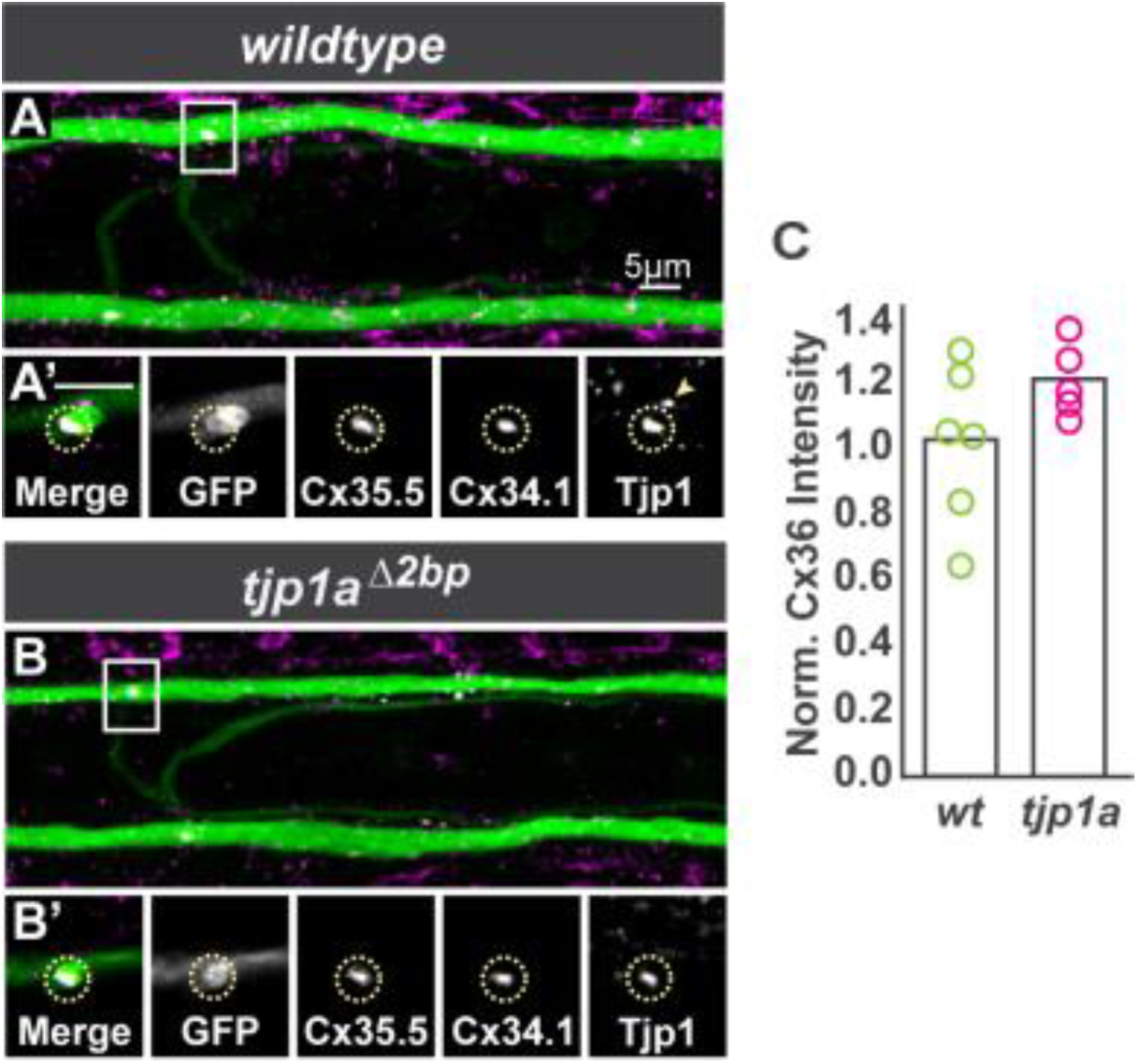
Electrical synapse formation does not require *tjp1a*. **A,B.** Images are dorsal view projections of ~5 um of a spinal cord segment from *M/CoLo:GFP* larvae at 5 days post fertilization. Larvae are stained for anti-GFP (green), anti-zebrafish-Cx35.5 (yellow), anti-zebrafish-Cx34.1 (cyan), and anti-human Tjp1/ZO1 (magenta). Boxes denote location of zooms. **A’,B’.** Individual Z-plane zooms of indicated region. Yellow circle denotes location of M/CoLo site of contact. Individual channels are shown in neighboring panels. In *tjp1a* mutants synaptic Cx and Tjp staining is unaffected (circles) but adjacent non-synaptic staining (arrowhead in **A’**) is lost in the mutants (**B’**). **C.** Bar graphs represent the mean of Cx36 quantified at synapses with each circle representing the average of 11-16 M/CoLo synapses within an animal.

**Fig. S3.**
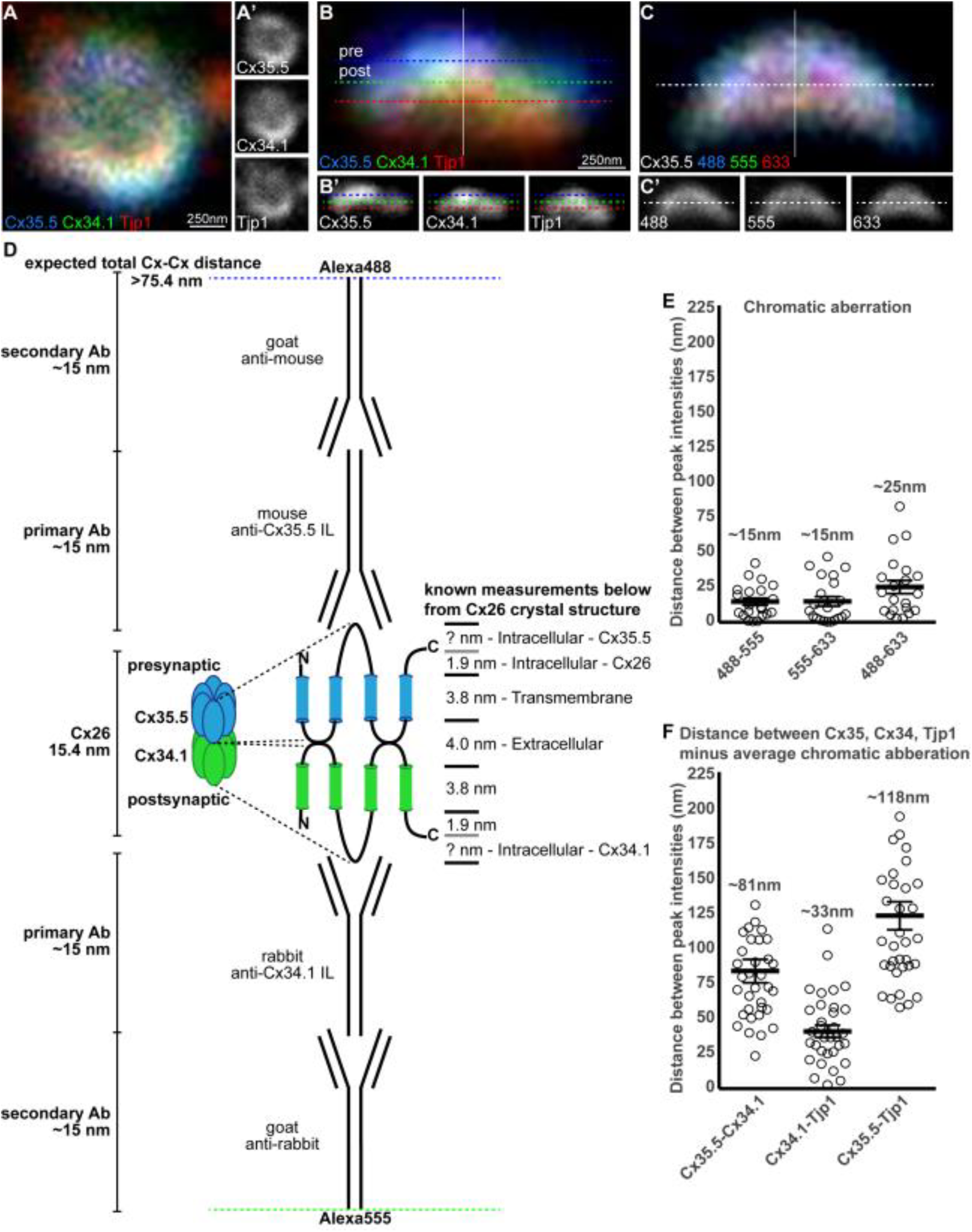
Analysis of presynaptic Cx35.5, postsynaptic Cx34.1, and Tjp1 staining at electrical synapses. **A-C.** Single plane magnifications of wildtype M/CoLo synapse showing *en face* (**A**) and side (**B**) orientations observed in the spinal cord. In (**A,B**) larvae are stained for anti-zebrafish-Cx35.5 (blue, stained with Alexa488), anti-zebrafish-Cx34.1 (green, stained with Alexa555), and anti-human Tjp1 (red, stained with Alexa633). In (**A**), the presynaptic Mauthner and postsynaptic CoLo are oriented into the page providing an *en face* view of the synapse, while in (**B**) Mauthner is at the top and the postsynaptic CoLo at the bottom providing a side view (neurons not shown). Individual channels are shown in neighboring panels. Our localization analysis focused on synapses that were in the side view (**B**) as these maximized the resolution for observing the location of staining. **C.** To measure the chromatic aberration we stained Cx35.5 with secondary antibodies in each dye (Alexa488, 555, and 633) used in localizing each protein of the synapse in (**A,B**). In (**B,C**) each dotted line represents the peak of fluorescence intensity for an individual fluorescent channel for the white line running vertically down the middle of the synapse. **D.** Theoretical Cx-Cx distance expected based on estimated sizes of Cx and antibodies. Known Cx distances are taken from the crystal structure of Cx26^35^. Note that Cx35.5 and Cx34.1 have longer intracellular loops than Cx26. The Cx35.5 and the Cx34.1 specific antibodies were made against the intracellular loops of each protein^9^. Antibody lengths are estimated from standard IgG proteins^36^. **E.** Estimation of chromatic aberration. Absolute distances between the peak fluorescent intensities of the noted channels from the Cx35.5 as stained in (**C**). The average measured chromatic aberration is noted. n = 21 individual synapses from 5 animals. **F.** Distance between the peak fluorescent intensities of the noted proteins as stained in (**B**). For each, the average chromatic aberration was removed to estimate the distance (Cx35.5-Cx34.1/488-555, Cx34.1-Tjp1/555-633, Cx35.5-Tjp1/488-633). The average measured distance between peak staining, minus the chromatic aberration, is noted. n = 35 individual synapses in 8 animals. For (**E,F**) each point represents the average distance at each individual synapse, with the line representing the average and the bars are standard error. Similar estimations of pre- and postsynaptic localizations have been applied to chemical synapses^37^.

## Extended Data

**Figure 1 Associated Data.**
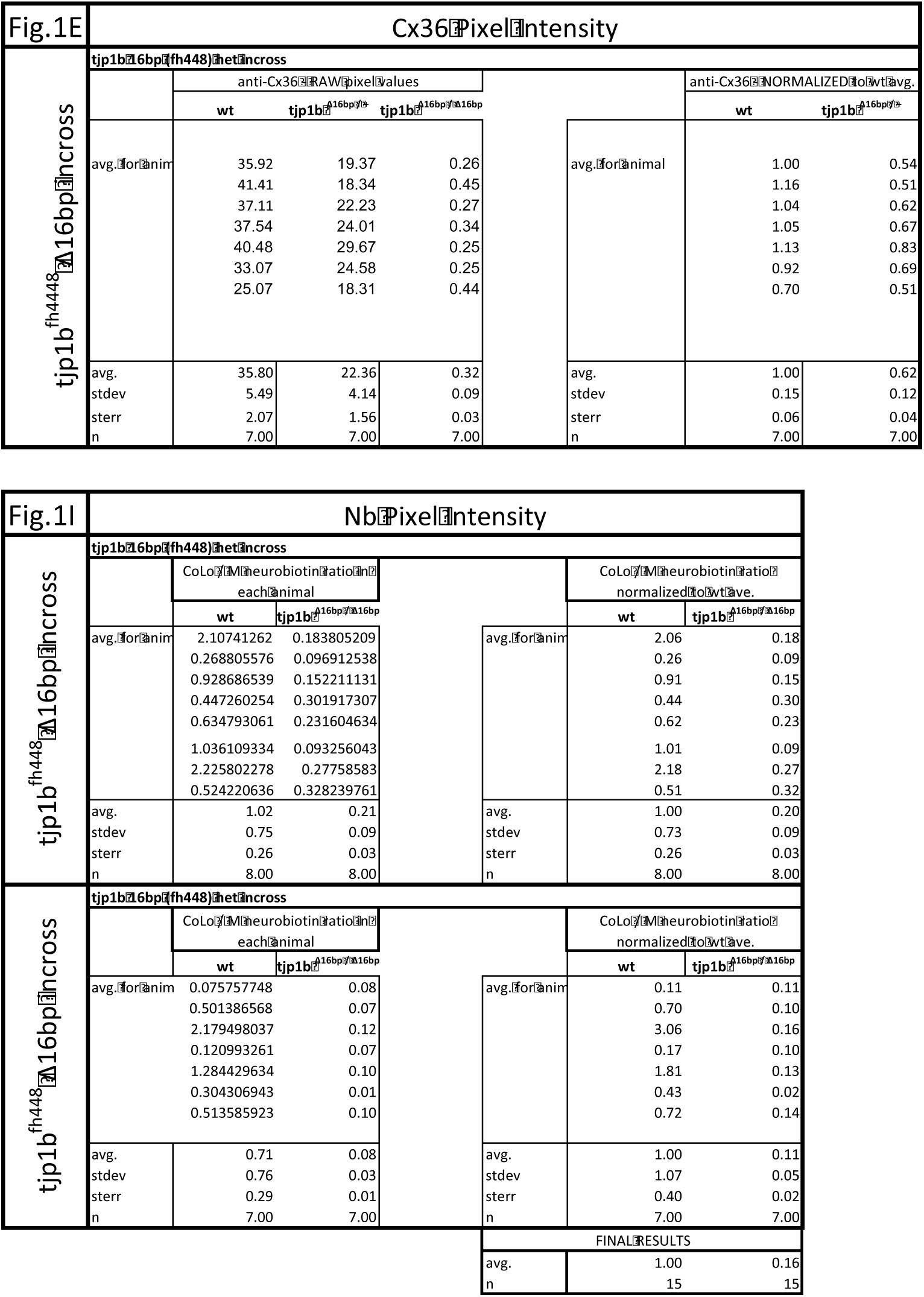

**Figure 2 Associated Data.**
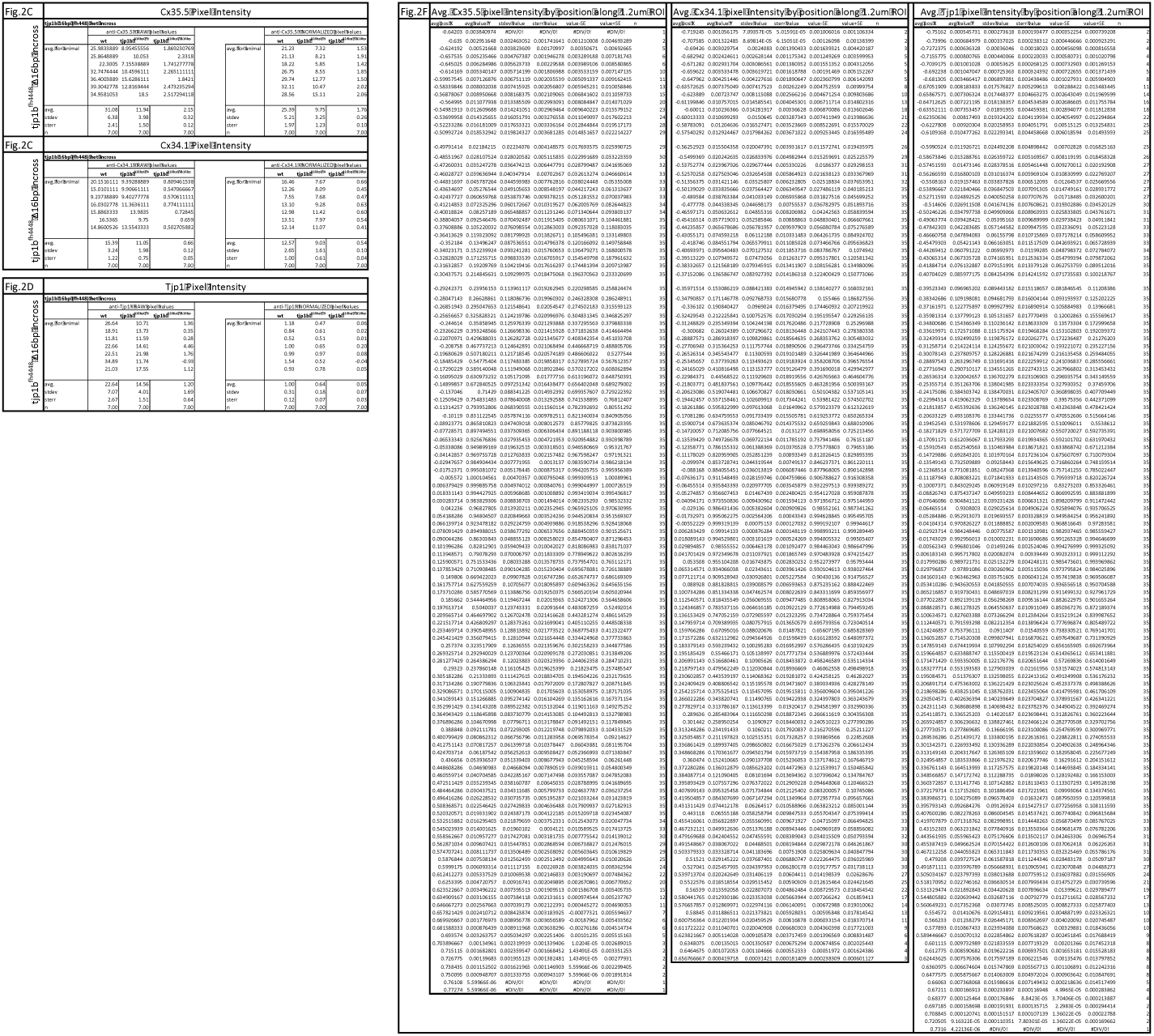

**Figure 3 Associated Data.**
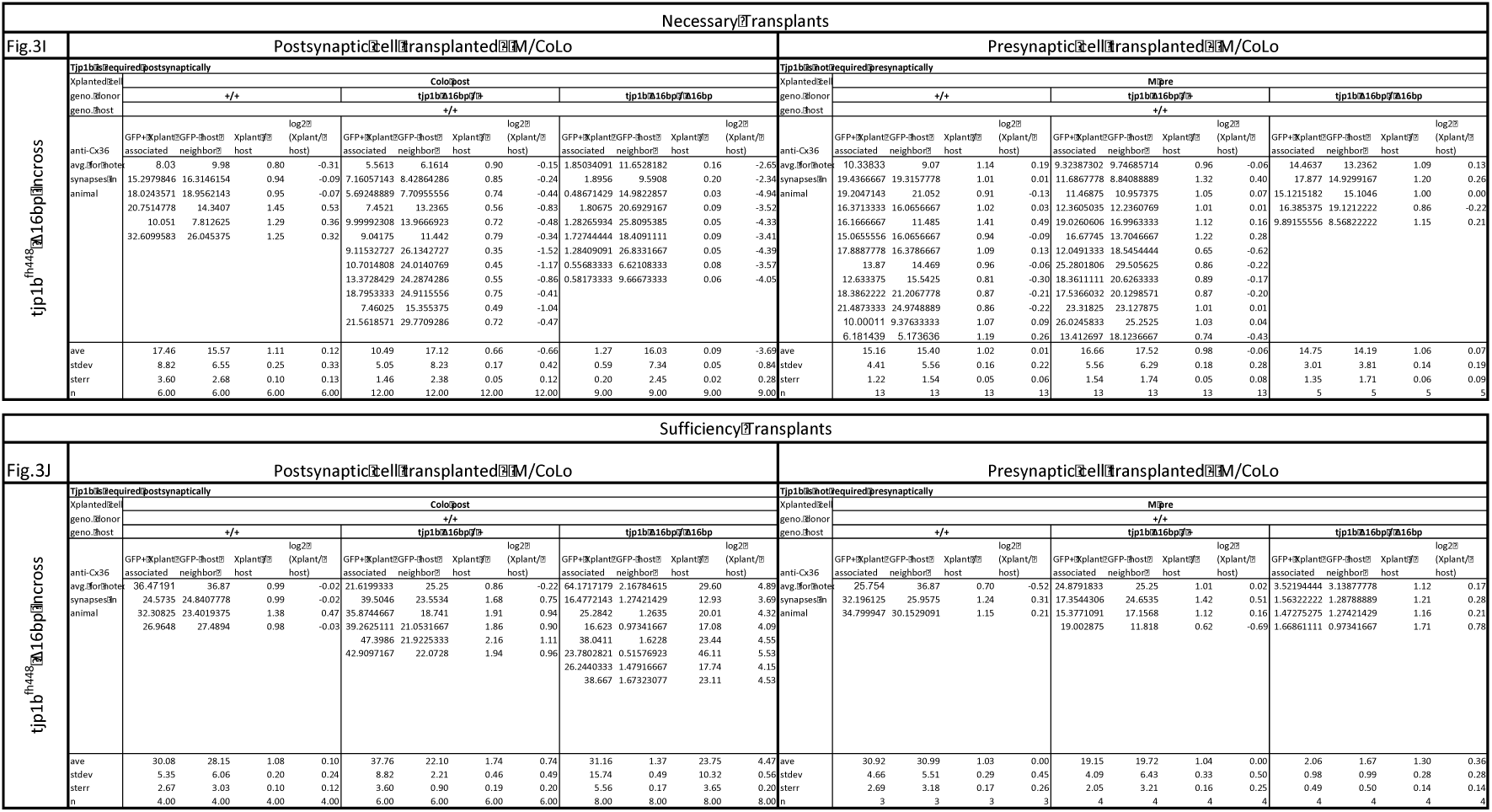

**Figure S1 Associated Data.**
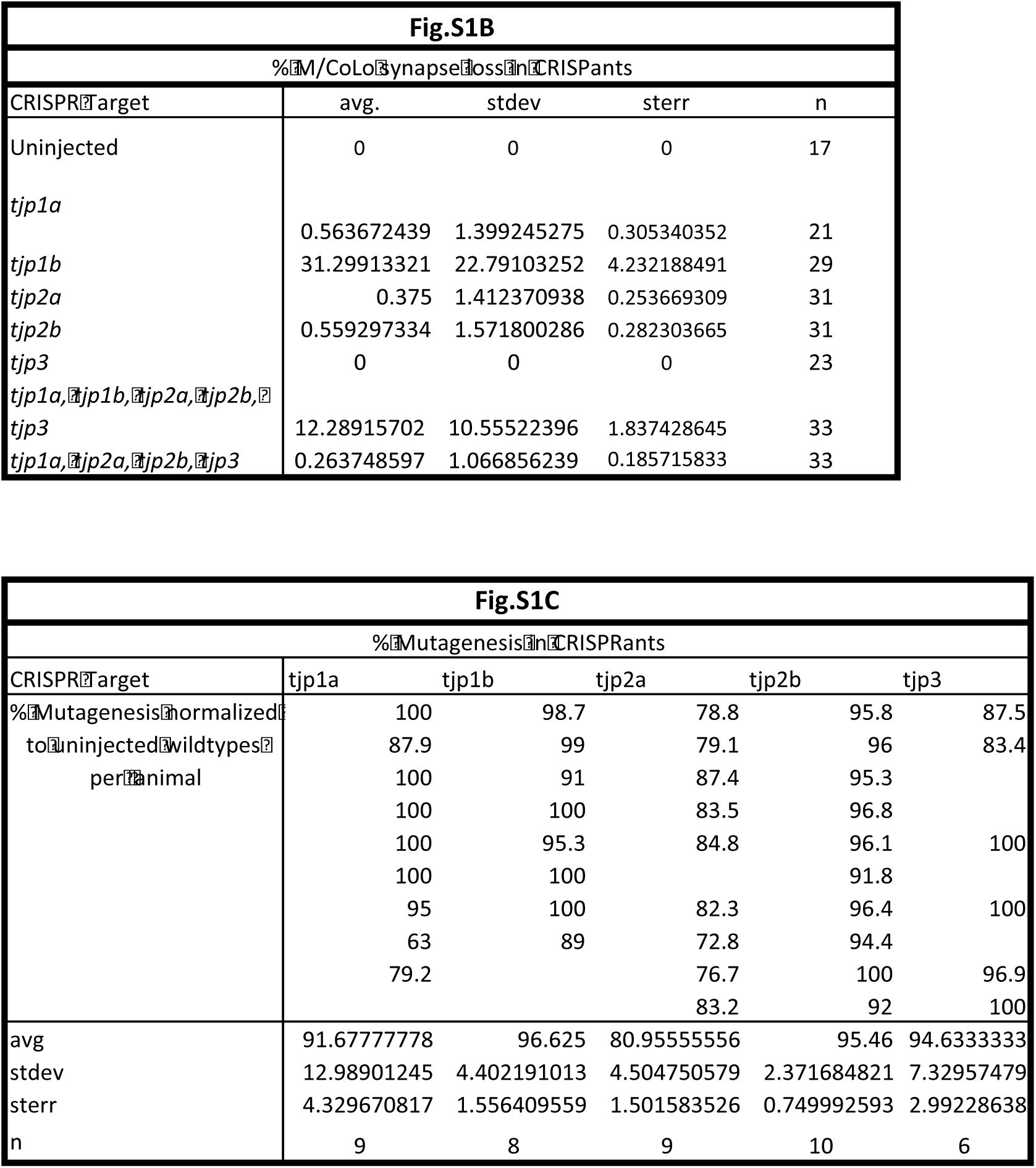

**Figure S2 Associated Data.**
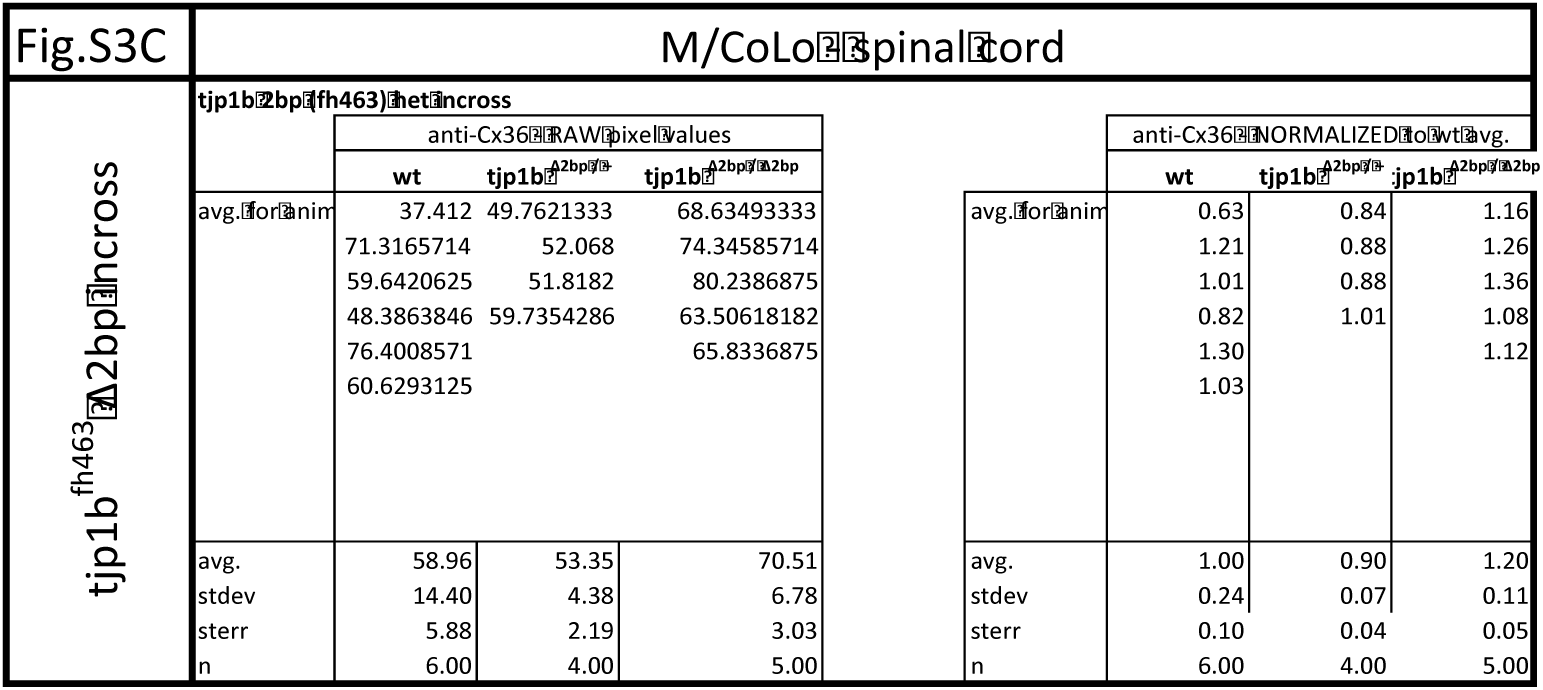

**Figure S3 Associated Data.**
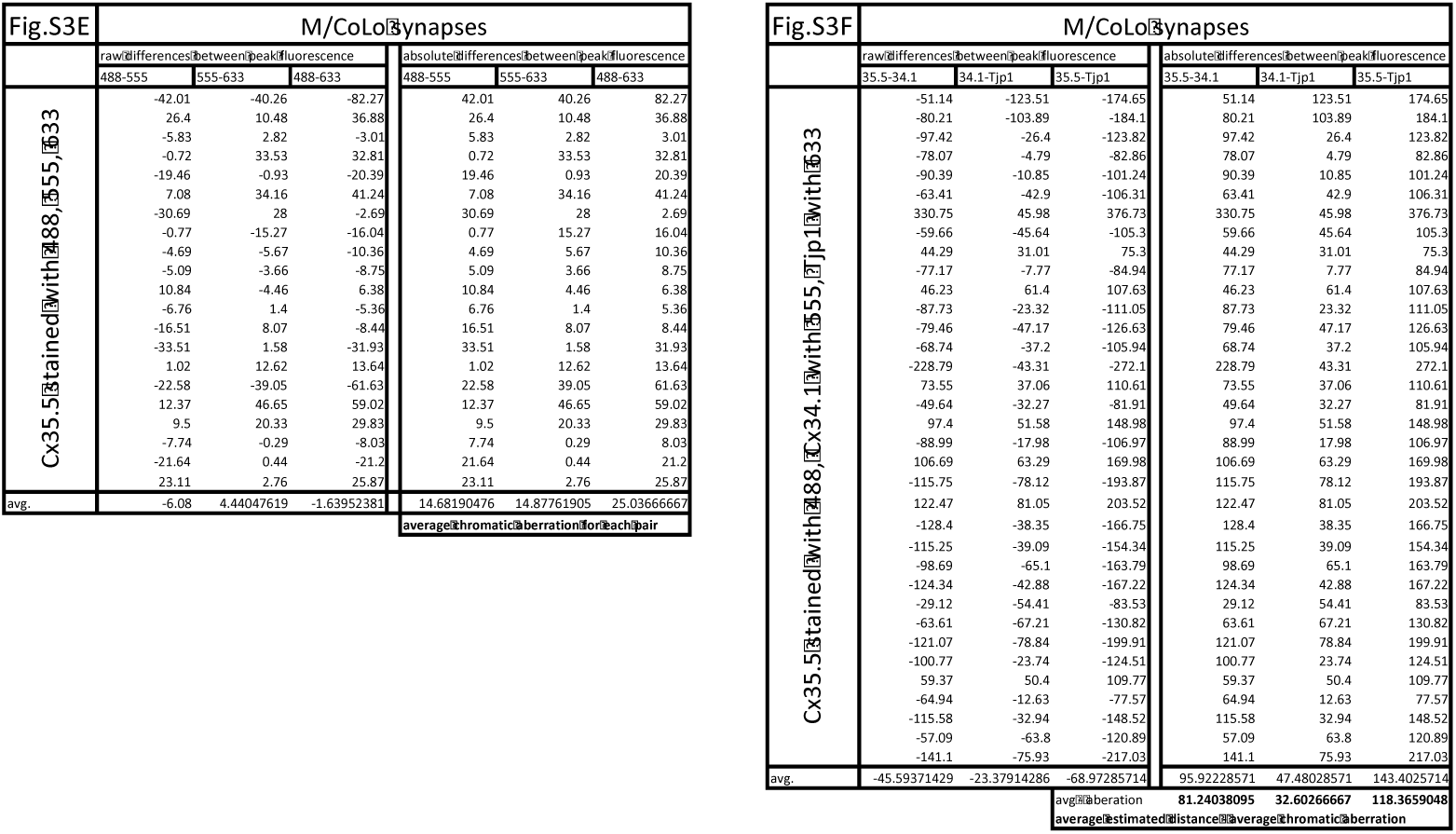

## Notes

**Support**: Funding was provided by the National Institute of Neurological Disease and Stroke R00NS085035 and startup funds from the University of Oregon to A.C.M.

We declare that we have no competing interests.

## References

1. Jin, Y. & Garner, C. C. Molecular Mechanisms of Presynaptic Differentiation. Annu. Rev. Cell Dev. Biol. 24, 237–262 (2008).

2. Zhu, J., Shang, Y. & Zhang, M. Mechanistic basis of MAGUK-organized complexes in synaptic development and signalling. Nat. Rev. Neurosci. 17, 209–223 (2016).

3. Connors, B. W. & Long, M. A. ELECTRICAL SYNAPSES IN THE MAMMALIAN BRAIN. Annu. Rev. Neurosci. 27, 393–418 (2004).

4. Pereda, A. E. Electrical synapses and their functional interactions with chemical synapses. Nat. Rev. Neurosci. 15, 250–263 (2014).

5. Phelan, P. et al. Molecular Mechanism of Rectification at Identified Electrical Synapses in the Drosophila Giant Fiber System. Curr. Biol. 18, 1955–1960 (2008).

6. Rash, J. E. et al. Molecular and Functional Asymmetry at a Vertebrate Electrical Synapse. Neuron 79, 957–969 (2013).

7. Phelan, P. et al. Molecular mechanism of rectification at identified electrical synapses in the Drosophila giant fiber system. Curr Biol. 18, 1955–1960 (2008).

8. Starich, T. a, Xu, J., Skerrett, I. M., Nicholson, B. J. & Shaw, J. E. Interactions between innexins UNC-7 and UNC-9 mediate electrical synapse specificity in the Caenorhabditis elegans locomotory nervous system. Neural Dev. 4, 16 (2009).

9. Miller, A. C. et al. A genetic basis for molecular asymmetry at vertebrate electrical synapses. Elife 6, (2017).

10. Brightman, M. W. & Reese, T. S. Junctions between intimately apposed cell membranes in the vertebrate brain. J. Cell Biol. 40, 648–677 (1969).

11. Sotelo, C. & Korn, H. Morphological correlates of electrical and other interactions through low-resistance pathways between neurons of the vertebrate central nervous system. Int. Rev. Cytol. **VOL** 55, 67–107 (1978).

12. Miller, A. C. & Pereda, A. E. The electrical synapse: Molecular complexities at the gap and beyond. Dev. Neurobiol. 77, 562–574 (2017).

13. Liu, K. S. & Fetcho, J. R. Laser Ablations Reveal Functional Relationships of Segmental Hindbrain Neurons in Zebrafish. Neuron 23, 325–335 (1999).

14. Satou, C. et al. Functional role of a specialized class of spinal commissural inhibitory neurons during fast escapes in zebrafish. J. Neurosci. 29, 6780–6793 (2009).

15. Miller, A. C., Voelker, L. H., Shah, A. N. & Moens, C. B. Neurobeachin is required postsynaptically for electrical and chemical synapse formation. Curr. Biol. 25, 16–28 (2015).

16. Shah, A. N., Davey, C. F., Whitebirch, A. C., Miller, A. C. & Moens, C. B. Rapid reverse genetic screening using CRISPR in zebrafish. Nat. Methods 12, 535–40 (2015).

17. Giepmans, B. N. & Moolenaar, W. H. The gap junction protein connexin43 interacts with the second PDZ domain of the zona occludens-1 protein. Curr. Biol. 8, 931–4

18. Balda, M. S. & Matter, K. Tight junctions as regulators of tissue remodelling. Curr. Opin. Cell Biol. 42, 94–101 (2016).

19. Li, X. et al. Neuronal connexin36 association with zonula occludens-1 protein (ZO-1) in mouse brain and interaction with the first PDZ domain of ZO-1. Eur. J. Neurosci. 19, 2132–2146 (2004).

20. Flores, C. E., Li, X., Bennett, M. V. L., Nagy, J. I. & Pereda, A. E. Interaction between connexin35 and zonula occludens-1 and its potential role in the regulation of electrical synapses. Proc. Natl. Acad. Sci. U. S. A. 105, 12545–12550 (2008).

21. Yao, C. et al. Electrical synaptic transmission in developing zebrafish. J. Neurophysiol. 552827, 2102–13 (2014).

22. Kiener, T. K., Sleptsova-Friedrich, I. & Hunziker, W. Identification, tissue distribution and developmental expression of tjp1/zo-1, tjp2/zo-2 and tjp3/zo-3 in the zebrafish, Danio rerio. Gene Expr. Patterns 7, 767–776 (2007).

23. O’Brien, J. The ever-changing electrical synapse. Current Opinion in Neurobiology 29, (2014).

24. Palacios-Prado, N., Huetteroth, W. & Pereda, A. E. Hemichannel composition and electrical synaptic transmission: molecular diversity and its implications for electrical rectification. Front. Cell. Neurosci. 8, 324 (2014).

25. Wang, X. et al. Neurobeachin: A protein kinase A-anchoring, beige/Chediak-higashi protein homolog implicated in neuronal membrane traffic. J. Neurosci. 20, 8551–8565 (2000).

26. Nair, R. et al. Neurobeachin regulates neurotransmitter receptor trafficking to synapses. J. Cell Biol. 200, 61–80 (2013).

27. Kimmel, C. B., Ballard, W. W., Kimmel, S. R., Ullmann, B. & Schilling, T. F. Stages of embryonic development of the zebrafish. Dev. Dyn. 203, 253–310 (1995).

28. Shah, A. N., Moens, C. B. & Miller, A. C. in Methods in cell biology 135, 89–106 (2016).

29. Schindelin, J. et al. Fiji: an open-source platform for biological-image analysis. Nat. Methods 9, 676–682 (2012).

30. Moreno-Mateos, M. A. et al. CRISPRscan: designing highly efficient sgRNAs for CRISPR-Cas9 targeting in vivo. Nat. Methods 12, 982–988 (2015).

31. Burger, A. et al. Maximizing mutagenesis with solubilized CRISPR-Cas9 ribonucleoprotein complexes. Development 143, (2016).

32. Carrington, B., Varshney, G. K., Burgess, S. M. & Sood, R. CRISPR-STAT: an easy and reliable PCR-based method to evaluate target-specific sgRNA activity. Nucleic Acids Res. 43, e157 (2015).

33. Kemp, H. A., Carmany-Rampey, A. & Moens, C. Generating Chimeric Zebrafish Embryos by Transplantation. J. Vis. Exp. (2009).

34. Chen, J., Pan, L., Wei, Z., Zhao, Y. & Zhang, M. Domain-swapped dimerization of ZO-1 PDZ2 generates specific and regulatory connexin43-binding sites. EMBO J. 27, 2113–2123 (2008).

35. Maeda, S. et al. Structure of the connexin 26 gap junction channel at 3.5 Å resolution. Nature 458, 597–602 (2009).

36. Wildanger, D., Medda, R., Kastrup, L. & Hell, S. W. A compact STED microscope providing 3D nanoscale resolution. J. Microsc. 236, 35–43 (2009).

37. Fouquet, W. et al. Maturation of active zone assembly by Drosophila Bruchpilot. J. Cell Biol. 186, (2009).

